# Characterization of the genome editing with miniature nucleases TnpB, IscB and enIscB in *Escherichia coli* strains

**DOI:** 10.1101/2024.09.04.611128

**Authors:** Hongjie Tang, Jie Gao, Mingjun Sun, Suyi Zhang, Qi Li

## Abstract

DNA nucleases TnpB and IscB were regarded as new antibacterial strategy to combat the drug-resistant bacteria represented by *Escherichia coli* due to its specificity in targeting DNA and smallest size, but the genome-editing of TnpB/IscB in *E. coli* remains unclear. This study characterized the genome-editing of TnpB/IscB in *E. coli* strains. First, the toxicity and cleavage results showed TnpB only worked in *E. coli* MG1655, while IscB and enIscB could perform in ATCC9637 and BL21(DE3). Next, TnpB-based genome-editing tool was established in MG1655, while IscB/enIscB achieved in ATCC9637/BL21(DE3). The copy number of TnpB/IscB/enIscB were changed to explore the impact of editing efficiency. Moreover, the editing plasmids were successfully cured. Finally, the escaping mechanism of *E. coli* under editing of TnpB/IscB was revealed. Overall, this study successfully applied TnpB/IscB/enIscB to genome-editing in *E. coli*, which will broaden genetic manipulation toolbox in *E. coli* and facilitate the development of new antimicrobial drugs.

## Introduction

The emergence of microbial drug resistance has intensified into a global concern due to its impact on health-care systems. *Escherichia coli*represents one of the most typical drug-resistant microorganisms that necessitate the urgent development of new antimicrobial drugs to combat it^1–7^. Although the development of novel antibiotics belongs to a prevalent strategy, this approach fails to avoid the novel drug resistance problems^8–11^. The RNA-guided DNA nucleases have been utilized as a new strategy for eliminating drug-resistant microorganisms according to its high specificity towards targeting DNA and efficient genome editing capabilities^12–15^. CRISPR-Cas9 (Clustered regularly interspaced short palindromic repeats, CRISPR; CRISPR associated nuclease, Cas9) has been reported to be used for eliminating of specific strains^16–21^, but the size of Cas9 is too large (∼1053-1368 amino acids, aa) to be effectively delivered to target cells^14^. Therefore, it is urgent to mine smaller RNA-guided DNA nucleases for eliminating drug-resistant microorganisms, which will accelerate the development of new antimicrobial drugs.

Recent research reported two smaller-sized RNA-guided DNA nucleases, including the TnpB (Transposon-associated transposase B, ∼400 aa) and the IscB (Insertion sequences Cas9-like B, IscB, ∼500 aa) encoded by the IS200/IS605 superfamily transposons, which was hypothesized to be the ancestors of Cas12 and Cas9 respectively^22,23^. The research findings on TnpB were as follows: it is discovered that TnpB protein encoded by the prokaryotic IS200/IS605 transposons could cleave double-strand DNA (dsDNA) under the guidance of the RNA, and TnpB was harnessed for genome editing in human cells^24,25^; the phylogenetic relationships of TnpB in six bacterial species were revealed by utilzing the genomes of high assembly levels (chromosome and complete)^26^; the mechanism of how TnpB recognized and cleaved its target DNA was subsequently elucidated by analyzing the detailed structure of the TnpB-ωRNA-dsDNA complex^27,28^; five hyper-compat editors within human cells were identified through the establishment of high-throughput screening of TnpB^29^; moreover, TnpB derived from *Sulfolobus islandicus* was established as genome editing in *Pediococcus acidilactici* and *Sulfolobus islandicus*^30^; it was reported that TnpB could optimized its own mRNA into ωRNA successfully^31^; recently, it is particularly noteworthy that precise genome editing and phenotypic rectification in a tyrosinaemia mouse model could be achieved by harnessing a transposon-linked TnpB-ωRNA system^32^; furthermore, a thermophilic archaeal TnpB with flexible TAM requirements was reprted, and it could improve the genome editing efficiency in the natural host^33^; the recent research found that TnpB obtained from *Firmicutes bacterium* could also performed dsDNA cleavage activity^34^.

For the research of IscB were as follows: the architectural similarities between IscB and Cas9 were demonstrated based on the structural analysis of IscB-ωRNA combined with a dsDNA target^35^; the process of how small IscB protein assembled with the ωRNA was revealed by analyzing the detailed architecture of the IscB-ωRNA complex^22,25^; additionally, a highly effcient of enIscB system in mammalian systems was obtained through optimizing the structure of wild-type IscB and ωRNA^36^.

The above reports illustrated the capability of two RNA-guided DNA nucleases for the applications of genome editing. The current research on TnpB, IscB and enIscB are mostly focused on genome editing in eukaryotic cells, while only one article reported that the genome editing can be achieved by TnpB in bacteria, and the application of TnpB, IscB and enIscB systems for genome editing in *E. coli* has not yet been reported. Therefore, in this study, *E. coli* MG1655, ATCC9637 and BL21 (DE3) were selected as the research objects to demonstrate the toxicity of TnpB, IscB and enIscB (enhanced IscB) proteins in those three *E. coli* strains. Subsequently, the genomic cleavage efficiencies of these three proteins in MG1655, ATCC9637 and BL21(DE3) were tested respectively. Furthermore, the gene deletion efficiencies of these three proteins in the above three strains were verified. And the effect of different plasmid copy numbers of TnpB, IscB and enIscB was also explored. Simultaneously, the plasmids curing of the TnpB/IscB/enIscB-based genome editing systems were achieved by two-step plasmid curing approach successfully. Finally, we also preliminarily explored the mechanisms of *E. coli* escaping under the editing of TnpB/IscB and revealed the main reason for the generation of escapers.

In summary, this study took the lead to characterize the application of TnpB, IscB and enIscB in *E. coli* strains including MG1655, ATCC9637 and BL21(DE3). Compared with Cas9 or Cas12a, the employment of smaller TnpB or IscB confers advantages in terms of vector construction and future delivery, will accelerate the development of new antimicrobial drugs based on the RNA-guided DNA nucleases.

## Results

### TnpB-based genome editing tools only worked in MG1655 and didn’t work in ATCC9637 and BL21(DE3)

As TnpB is currently the smallest programmable RNA-guided DNA nuclease, the *Sulfolobus islandicus*-derived TnpB was firstly focused to explore the genome editing efficiency in *E. coli*^30^. First, the plasmid that the p15A-TnpB and the pCK1 plasmid (control group) were transformed into *E. coli* MG1655, ATCC9637 or BL21(DE3) respectively (Fig. 1A), and the toxicity of TnpB protein in *E. coli* was determined by counting the transformation efficiency. The results demonstrated that the transformation efficiencies of p15A-TnpB in MG1655 and BL21(DE3) were similar to pCK1 plasmid (106 CFU/μg plasmid DNA), indicating that TnpB protein was nontoxic to MG1655 and BL21(DE3) (Fig. 1B, C). Unfortunately, a significant reduction of transformation efficiencies of p15A-TnpB were observed in ATCC9637, which was approximately 102 CFU/μg plasmid DNA (Fig. 1B). Even if the copy number of TnpB was reduced, the transformation efficiency of pSC101-TnpB in ATCC9637 was still not improved (Supplementary Fig. 1A, B). These above results suggested that the TnpB was toxic to ATCC9637 and hindered the application of TnpB for genome editing in this host. Thus, the further testing of cleavage activities of TnpB was carried out only in MG1655 and BL21(DE3).

**Fig. 1.**
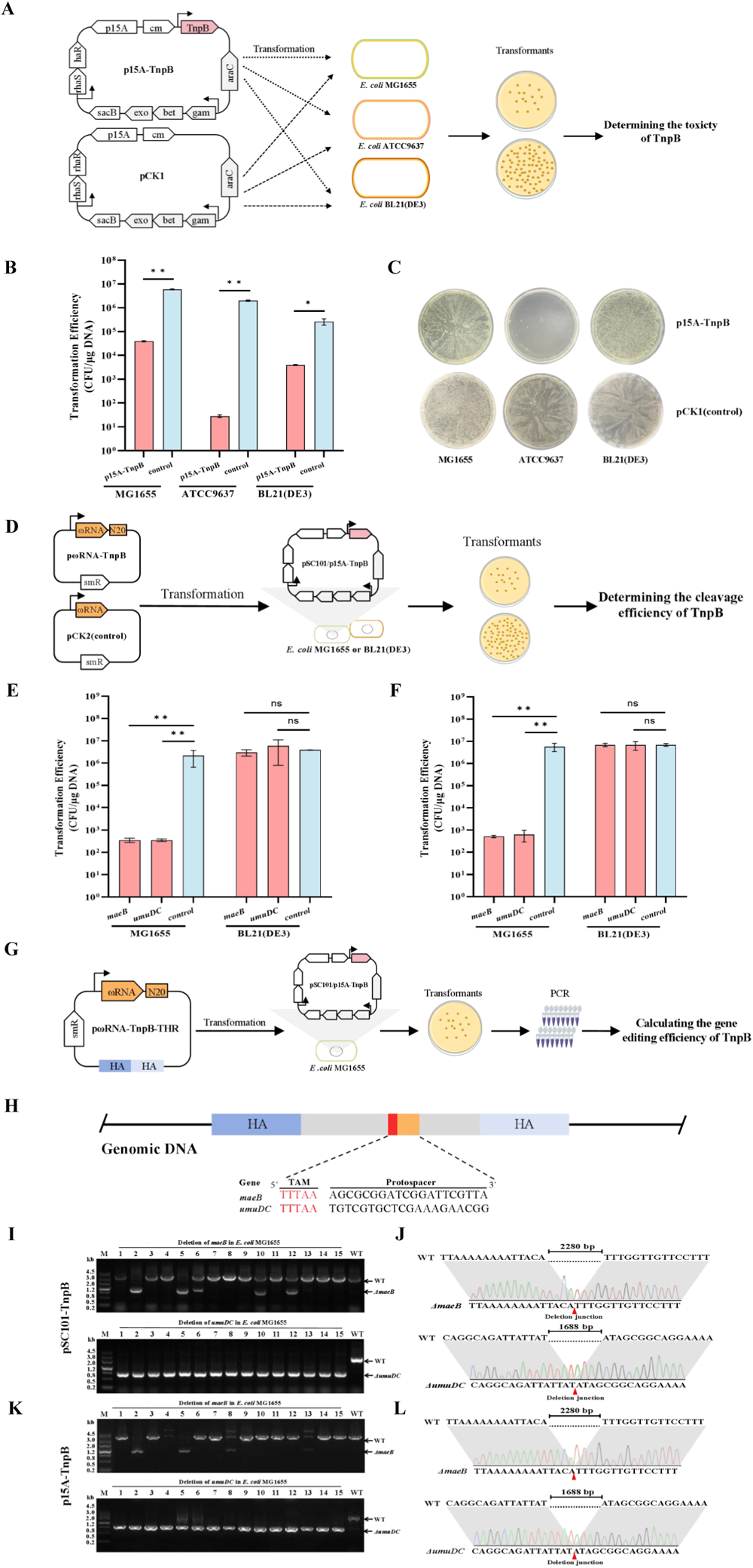
TnpB-based genome editing tool was successfully established in *E. coli* MG1655, but not in *E. coli* ATCC9637 and BL21(DE3). **A** The detailed diagram of the testing of toxicity of p15A-TnpB in *E. coli* MG1655, ATCC9637 and BL21(DE3). **B** The calculation of toxicity of p15A-TnpB in *E. coli* MG1655, ATCC9637 and BL21(DE3). **C** The plating results for the verification of toxicity of p15A-TnpB in *E. coli* MG1655, ATCC9637 and BL21(DE3). **D** The workflow diagram for testing the cleavage efficiencies of pSC101-TnpB/p15A-TnpB in *E. coli* MG1655 and BL21(DE3). **E** The genomic cleavage efficiencies in *E. coli* MG1655 and BL21(DE3) by using pSC101-TnpB. **F** The genomic cleavage efficiencies in *E. coli* MG1655 and BL21(DE3) by using p15A-TnpB. **G** The workflow for gene deletion assay with pSC101-TnpB or p15A-TnpB in *E. coli* MG1655. HA means homology arm. **H** The targeted sites for gene deletion in *E. coli* MG1655. **I** PCR amplification results of the deletion of *maeB* and *umuDC* genes in *E. coli* MG1655 by using pSC101-TnpB. **J** DNA sequencing results of the deletion of *maeB* and *umuDC* genes in *E. coli* MG1655 by using pSC101-TnpB. **K** PCR amplification results of the deletion of *maeB* and *umuDC* genes in *E. coli* MG1655 by using p15A-TnpB. **L** DNA sequencing results of the deletion of *maeB* and *umuDC* genes in MG1655 by using pSC101-TnpB. ns, not significant; *P < 0.05; **P < 0.01.

The RNA-guided DNA nucleases cleave bacterial genomic DNA to form a double-strand break (DSB), and the DSB is usually repaired by homologous recombination or nonhomologous end joining (NHEJ) pathways^37–39^. Only a few bacteria contain the NHEJ^40^. Hence, most of the current genome editing tools in bacteria are depended on providing the homologous arms exogenously to repair the DSBs by homologous recombination^41,42^. In the absence of an exogenous repair template, the DSBs in bacteria caused by DNA nucleases typically are irreparable, leading to cell death. Based on this principle, the cleavage activities of TnpB in MG1655, BL21(DE3) were therefore tested without the provision of repair homologous arms. The pωRNA-TnpB-*maeB* or pωRNA-TnpB-*umuDC* plasmid that targeting *maeB* or *umuDC* gene and the pCK2 plasmid (control group) was respectively transferred into MG1655, BL21(DE3) carrying the pSC101-TnpB or p15A-TnpB (Fig. 1D). It was noteworthy that the transformation efficiency of pωRNA-TnpB-*maeB* or pωRNA-TnpB-*umuDC* in MG1655 harboring pSC101-TnpB or p15A-TnpB was approximately 10^2^ CFU/μg plasmid DNA, which was dropped 4 orders of magnitude compared to the control group. While in BL21(DE3), the transformation efficiencies were similar to the control plasmid, with approximately 10^6^ CFU/μg plasmid DNA (Fig. 1E, F and Supplementary Fig. 2A, B). The above findings suggested that TnpB performed the cleavage activity towards MG1655, but it failed to perform the cleavage activity in BL21(DE3). Consequently, the gene deletion efficiency of TnpB was measured only in MG1655.

To repair the DSB formed by the excision of TnpB in MG1655, the upstream and downstream repair homoloy arms (in the form of linear dsDNA) of *maeB* or *umuDC* gene, along with the corresponding pωRNA-TnpB-targetX plasmids, were co-electroporated into MG1655 carrying pSC101-TnpB or p15A-TnpB. A certain number of colonies could be obtained on the plates, and about 15 colonies were randomly selected for verifying by colony PCR. However, no positive colonies were obtained. It was speculated that the double-stranded form of homology arms in MG1655 carrying pSC101-TnpB or p15A-TnpB might be unstable that leading to its degradation. Thus, the homology arms were inserted into pωRNA-TnpB-*maeB* or pωRNA-TnpB-*umuDC* to generate plasmids pωRNA-TnpB-*maeB*-THR or pωRNA-TnpB-*umuDC*-THR respectively. Similarly, these plasmids were electroporated into MG1655 containing pSC101-TnpB or p15A-TnpB respectively (Fig. 1G, H and Table 1). Then about 15 colonies were randomly selected and verified by colony PCR. Fortunately, both *maeB* and *umuDC* genes were deleted successfully with efficiencies of 30% and 100%. The PCR products were further sequenced to verify the positive editing (Fig. 1I-L and Table 1). The above results showed that the gene deletion in *E. coli* MG1655 can be achieved by using the TnpB, but the homology arms need to be provided in the form of plasmids rather than linear dsDNA.

**Table 1.**
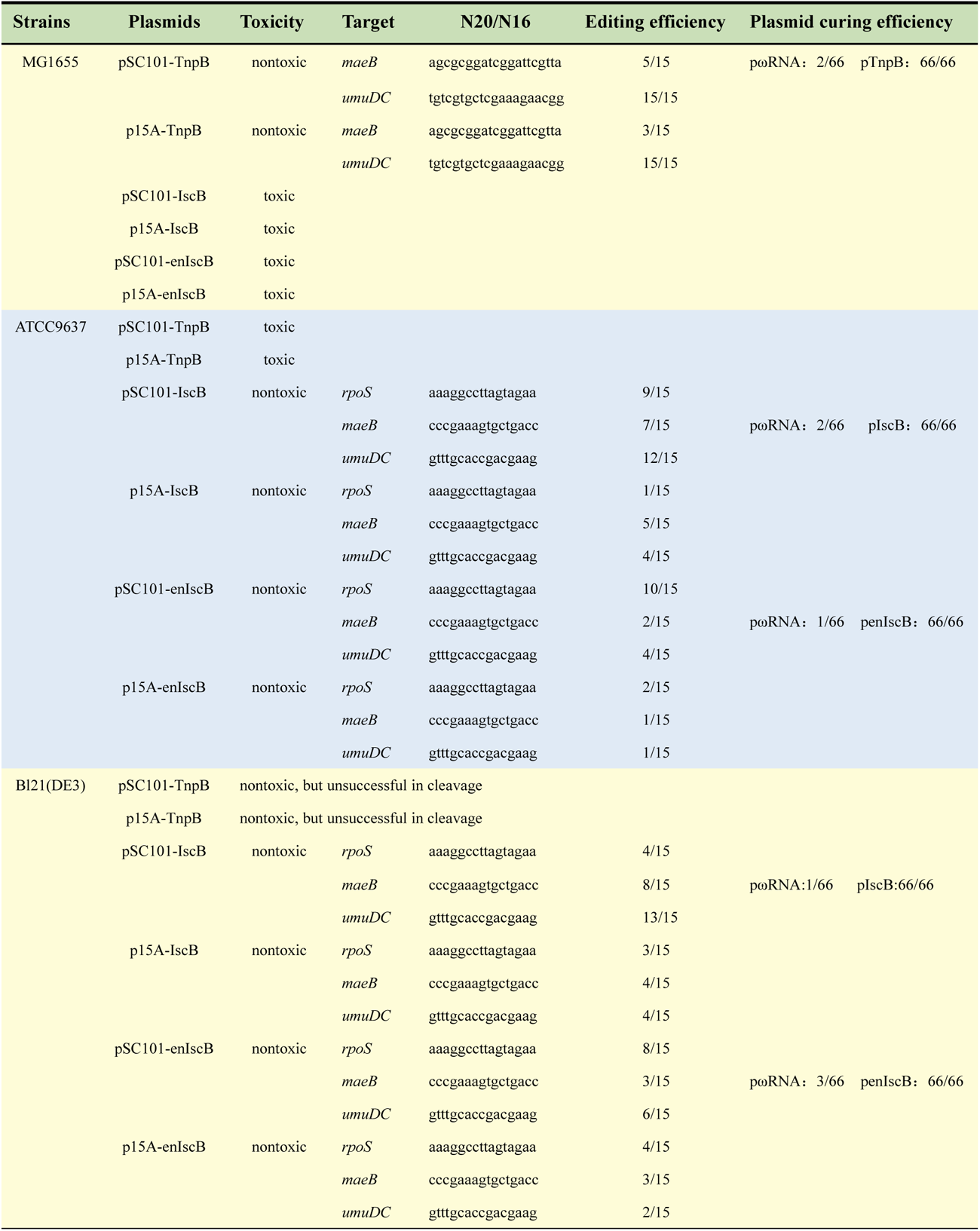
The summary of the application of the pTnpB/pωRNA, pIscB/pωRNA and penIscB/pωRNA systems for genome editing in *E. coli* MG1655, ATCC9637 and BL21(DE3).

### Efficient genome editing could be achieved in ATCC9637 and BL21 (DE3) by using IscB, but not in MG1655

The above results showed that TnpB-based genome editing can only be realized in MG1655, indicating the limitation of TnpB’s application. Therefore, our attention was moved to the IscB protein, which is derived from the human intestinal macro genome^36^, and it also has a small size (∼500 amino acids). We aimed to explore whether IscB could fill the gap of TnpB. First, the pSC101-IscB plasmid and the pCK1 plasmid (control group) were introduced into *E. coli* MG1655, ATCC9637 or BL21(DE3) respectively. After counting the transformation efficiency, the results showed that the transformation efficiencies of pSC101-IscB in ATCC9637 and BL21(DE3) were consistent to the control (∼10^6^ CFU/μg plasmid DNA), indicating that IscB protein was nontoxic toward ATCC9637 and BL21(DE3) (Fig. 2A). Whereas, the transformation efficiency in MG1655 was reduced about 4 orders of magnitude compared to the control plasmid, which indicated that IscB was toxic to MG1655(Fig. 2A and Supplementary Fig. 1C). Based on the above phenomena, the subsequent testing of the cleavage activities of IscB was carried out only in ATCC9637 and BL21(DE3).

**Fig. 2.**
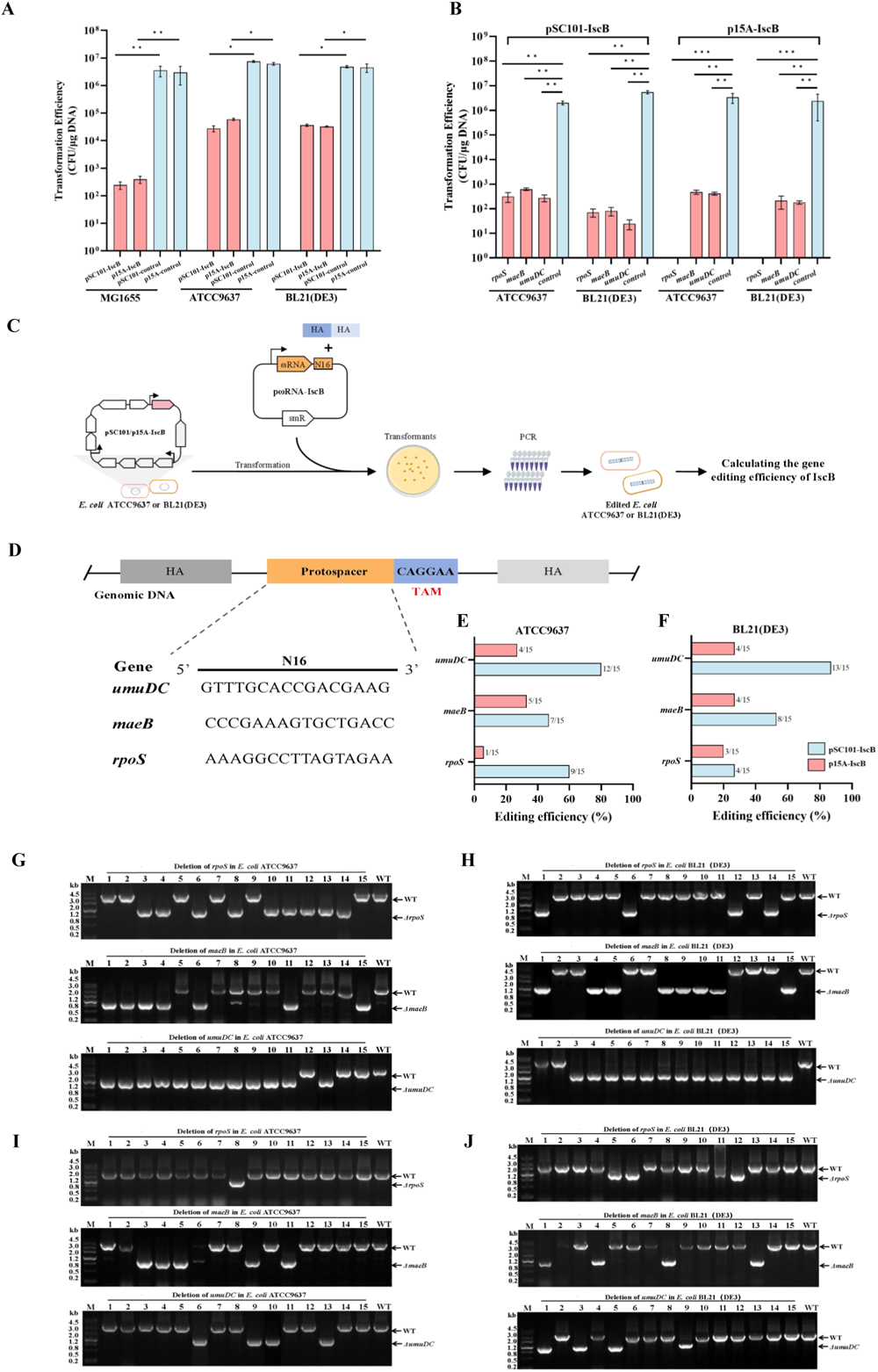
The genome editing could be achieved in *E. coli* ATCC9637 and BL21(DE3) while not in MG1655 by using IscB. **A** The verification results of toxicity of pSC101-IscB/p15A-IscB in *E. coli* MG1655, ATCC9637 and BL21(DE3). **B** The genome cleavage efficiencies in ATCC9637 and BL21(DE3) by using pSC101-IscB/p15A-IscB. **C** Schematic diagram of pSC101-IscB/p15A-IscB-mediated gene editing in *E. coli* ATCC9637 and BL21(DE3). **D** The targeted sites of the different genes in *E. coli* ATCC9637 and BL21(DE3). **E** Gene deletion efficiencies of pSC101-IscB or p15A-IscB in *E. coli* ATCC9637. **F** Gene deletion efficiencies of pSC101-IscB or p15A-IscB in *E. coli* BL21(DE3). **G** The colony PCR results of the deletion of *rpoS, maeB and umuDC* genes in ATCC9637 by using pSC101-IscB. **H** The colony PCR results of the deletion of *rpoS, maeB and umuDC* genes in BL21(DE3) by using pSC101-IscB. **I** PCR amplification results of the deletion of *rpoS, maeB and umuDC* genes in ATCC9637 by using p15A-IscB. **J** PCR amplification results of the deletion of *rpoS, maeB and umuDC* genes in BL21(DE3) by using p15A-IscB. ns, not significant; *P < 0.05; **P < 0.01; ***P < 0.001.

Similarly, the pωRNA-IscB-*rpoS,* pωRNA-IscB-*maeB* or pωRNA-IscB-*umuDC* plasmids that anchoring *rpoS, maeB or umuDC* genes and the pCK2 plasmid (control group) were electroporated into ATCC9637, BL21(DE3) harboring pSC101-IscB respectively. It was found that the transformation efficiencies of the pωRNA-IscB-*rpoS,* pωRNA-IscB-*maeB* or pωRNA-IscB-*umuDC* plasmids in ATCC9637 and BL21(DE3) harboring pSC101-IscB were approximately 10^2^ CFU/μg plasmid DNA, which was significantly lower than the control (∼10^6^ CFU/μg plasmid DNA) (Fig. 2B and Supplementary Fig. 3A). The above findings indicated that IscB protein could cleavage the ATCC9637 and BL21(DE3). Subsequently, the gene deletion efficiencies of IscB were further tested in ATCC9637 and BL21(DE3) on this basis.

The repair homology arms of *rpoS, maeB* or *umuDC* genes (in the linear dsDNA form) together with the corresponding pωRNA-IscB-targetX plasmids were co-electroporated into ATCC9637, BL21(DE3) containing pSC101-IscB to assess the gene deletion efficiency (Fig. 2C, D and Table 1). About 15 colonies were randomly picked for PCR verification. The results showed *maeB* and *umuDC* genes were deleted with efficiencies of 50% and 80% approximately in the above two *E. coli* strains, and the editing efficiency of *rpoS* gene ranged from 30%∼60% (Fig. 2E-H and Supplementary Fig. 4A, B). The above results showed that IscB could realize the genome editing in ATCC9637 and BL21(DE3).

Based on the above research, it was found that increasing the copy number of Cas9 could improve the genome editing efficiency in *E. coli*^43^. Thus, it was speculated that whether increasing the copy number of IscB could impact the genome editing efficiency in *E. coli*. IscB next was placed on p15A plasmid backbone which carrying a higher copy number (∼10 copies). Similarly, the p15A-IscB plasmid and the pCK1 (control group) plasmid were transformed into ATCC9637 and BL21(DE3) respectively to test the toxicity of IscB. The results showed that the transformation efficiencies of p15A-IscB in ATCC9637 and BL21(DE3) were approximately 106 CFU/μg plasmid DNA, which were similar to the control (Fig. 2A and Supplementary Fig. 1C). Then, the pωRNA-IscB-*rpoS,* pωRNA-IscB-*maeB* or pωRNA-IscB-*umuDC* plasmids that targeting *rpoS, maeB* or *umuDC* genes and the pCK2 (control) plasmid were respectively transformed into ATCC9637, BL21(DE3) containing p15A-IscB. It was found that all three genes could be cleaved by using p15A-IscB, in which the transformation efficiencies of both pωRNA-IscB-*maeB* and pωRNA-IscB-*umuDC* were approximately 10^2^ CFU/μg plasmid DNA. While for pωRNA-IscB-*rpoS,* was reduced 6 orders of magnitude compare to the control, with the transformation efficiency of 0 (Fig. 2B and Supplementary Fig. 3B). The above results showed that the cleavage efficiency of *rpoS* gene could be improved through increasing the copy number of IscB.

Furthermore, the repair homology arms of *rpoS, maeB* or *umuDC* genes along with the corresponding pωRNA-IscB-targetX plasmids were co-electroporated into ATCC9637, BL21(DE3) harboring p15A-IscB respectively (Fig. 2C and Table 1). About 15 colonies grown on the plates were picked randomly and checked by colony PCR. According to the results of DNA gel electrophoresis, the editing efficiencies at *maeB* and *umuDC* sites were approximately 30%, while the editing efficiency of *rpoS* site ranged from 6% to 20% in the above two *E. coli* (Fig. 2E, F, I, J and Supplementary Fig. 4C, D). Compared with the gene deletion efficiency of pSC101-IscB, the above findings illustrated that increasing the copy number of IscB has no significant effect on the improving of genome editing efficiency (Table 1).

### The genome editing in *E. coli* ATCC9637 and BL21(DE3) could be also realized by using enIscB

The previous study has performed several rounds of optimization of IscB and obtained an enIscB system (Fig. 3A) which could significantly improve base-editing efficiency in eukaryotic cells^36^. Therefore, the purpose of this part study was to explore whether enIscB could improve the genome editing efficiency in *E. coli*. First, the pSC101-enIscB plasmid and the pCK1 plasmid (control) were transformed into MG1655, ATCC9637 or BL21(DE3) respectively. It was found that the result of toxicity testing of enIscB in the three *E. coli* were similar to the IscB, the transformation efficiency of enIscB in MG1655 was 102 CFU/μg plasmid DNA, while in ATCC9637 and BL21(DE3) was 106 CFU/μg plasmid DNA (Fig. 3B and Supplementary Fig. 1D). Therefore, the subsequent testing of the cleavage activities of enIscB was carried out only in ATCC9637 and BL21(DE3). The pωRNA-IscB-*rpoS,* pωRNA-IscB-*maeB* or pωRNA-IscB-*umuDC* plasmid and the pCK2 (control) plasmid were respectively transformed into ATCC9637 or BL21(DE3) containing pSC101-enIscB. The results showed that the *rpoS, maeB* and *umuDC* genes could be cleaved in the above *E. coli* by utilizing enIscB, with the transformation efficiencies of 0, 10^2^ and 10^2^ CFU/μg plasmid DNA (Fig. 3C and Supplementary Fig. 5A).

**Fig. 3.**
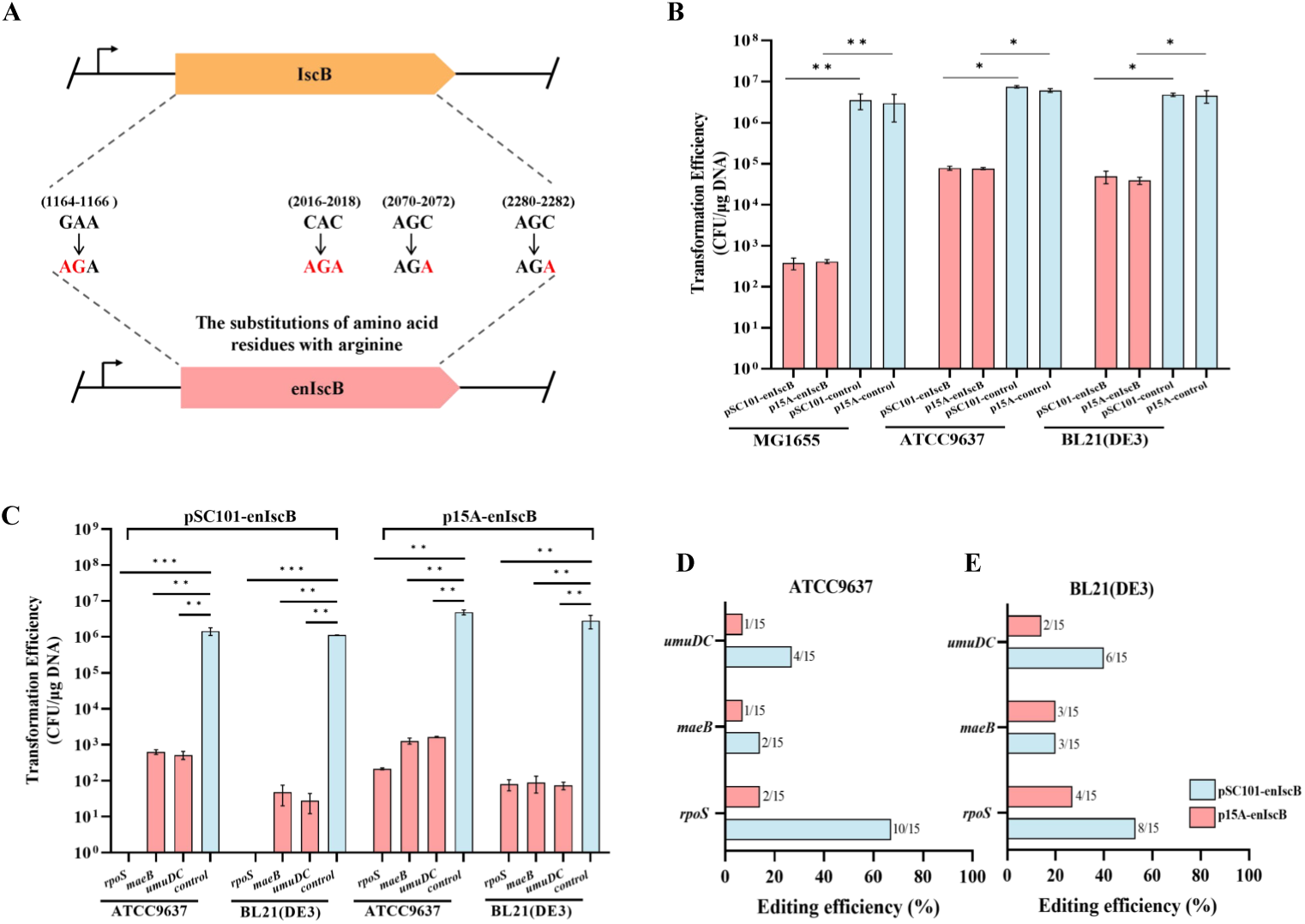
EnIscB-based genome editing tool could be established in *E. coli* ATCC9637 and BL21(DE3), but not in *E. coli* MG1655. **A** Schematic illustration of the enIscB variant gene structure. **B** The verification results of toxicity of pSC101-enIscB/p15A-enIscB in *E. coli* MG1655, ATCC9637 and BL21(DE3). **C** The cleavage efficiencies of pSC101-enIscB/p15A-enIscB at different targeted genes in *E. coli* ATCC9637 and BL21(DE3). **D** Gene deletion efficiencies in *E. coli* ATCC9637 by using pSC101-enIscB or p15A-enIscB. **E** Gene deletion efficiencies in *E. coli* BL21(DE3) by using pSC101-enIscB or p15A-enIscB. *P < 0.05; **P < 0.01; ***P < 0.001.

Next, the repair homology arms (also in the form of linear dsDNA) of *rpoS, maeB* or *umuDC* genes together with the corresponding pωRNA-IscB-targetX plasmids were co-electroporated into ATCC9637 or BL21(DE3) containing pSC101-enIscB respectively (Table 1). Following colony PCR, the sizes of target bands were analyzed. It was showed that these three genes were successfully edited, the editing efficiency of *rpoS* gene in ATCC9637 and BL21 (DE3) was 70%, while the editing efficiencies of *maeB* and *umuDC* genes varied from 10% to 40% (Fig. 3D, E and Supplementary Fig. 6A, B). The results showed that the enIscB system possessed a comparable editing efficiency to the IscB system.

Based on the above results, we further wondered the effect of increasing the copy number of enIscB on the genome editing efficiency in *E. coli*. Thus, the enIscB gene was subsequently inserted into the p15A plasmid vector. The methods for testing the toxicity, cleavage efficiencies and gene deletion efficiencies of p15A-enIscB in ATCC9637 and BL21(DE3) were kept the same as before increasing the copy number of enIscB (Table 1). It was found that the transformation efficiencies of pωRNA-IscB-*maeB* and pωRNA-IscB-*umuDC* were 102 CFU/μg plasmid DNA in the above *E. coli*, and the results were essentially consistent with that increasing the copy number of enIscB previously (Fig. 3C and Supplementary Fig. 5B). Next, gene deletion efficiencies of p15A-enIscB in ATCC9637 and BL21(DE3) were tested. About 15 colonies were randomly selected and checked, and it was found that the *rpoS, maeB* and *umuDC* genes were successfully deleted in ATCC9637 and BL21(DE3), with the editing efficiencies ranging from 6% to 30% (Fig. 3D, E and Supplementary Fig. 6C, D). The above results showed that enIscB-based genome editing tool in ATCC9637 and BL21(DE3) could also be realized by increasing the content of homology arm.

### The plasmids elimination of TnpB/IscB/enIscB-based genome editing systems were achieved by two-step plasmid curing strategy

Plasmids involved in genome editing should be eliminated for the further genome phenotypic testing or the next-round round of genome editing. A two-step plasmid curing strategy was devised, wherein pωRNA series were eliminated first and pTnpB/pIscB/penIscB were subsequently cured (Fig. 4A). Here, the *maeB* mutant of the *E. coli* MG1655 containing pSC101-TnpB was taken as an example. The edited MG1655 strain containing pSC101-TnpB was cultured overnight in liquid LB medium only contained kanamycin (50 µg/mL), and after two rounds of growth for approximately 24 h, single colonies were isolated and screened for sensitivity to spectinomycin (50 µg/mL) and resistance to kanamycin (50 µg/mL) by plating onto LB plates to confirm loss of pωRNA-TnpB-*maeB*-THR. Next, the pSC101-TnpB carried the levansucrase gene *sacB*, the product of *SacB* can convert sucrose to levan, which is highly toxic to *E. coli*^44^. Thus, the gene *sacB* was used as a counterselective marker. The edited MG1655 strain was then cultured and screened on LB plates containing sucrose (10 g/L). The positive colonies were further tested for sensitivity to kanamycin (50 µg/mL) to confirm loss of pSC101-TnpB plasmids (Fig. 4A).

**Fig. 4.**
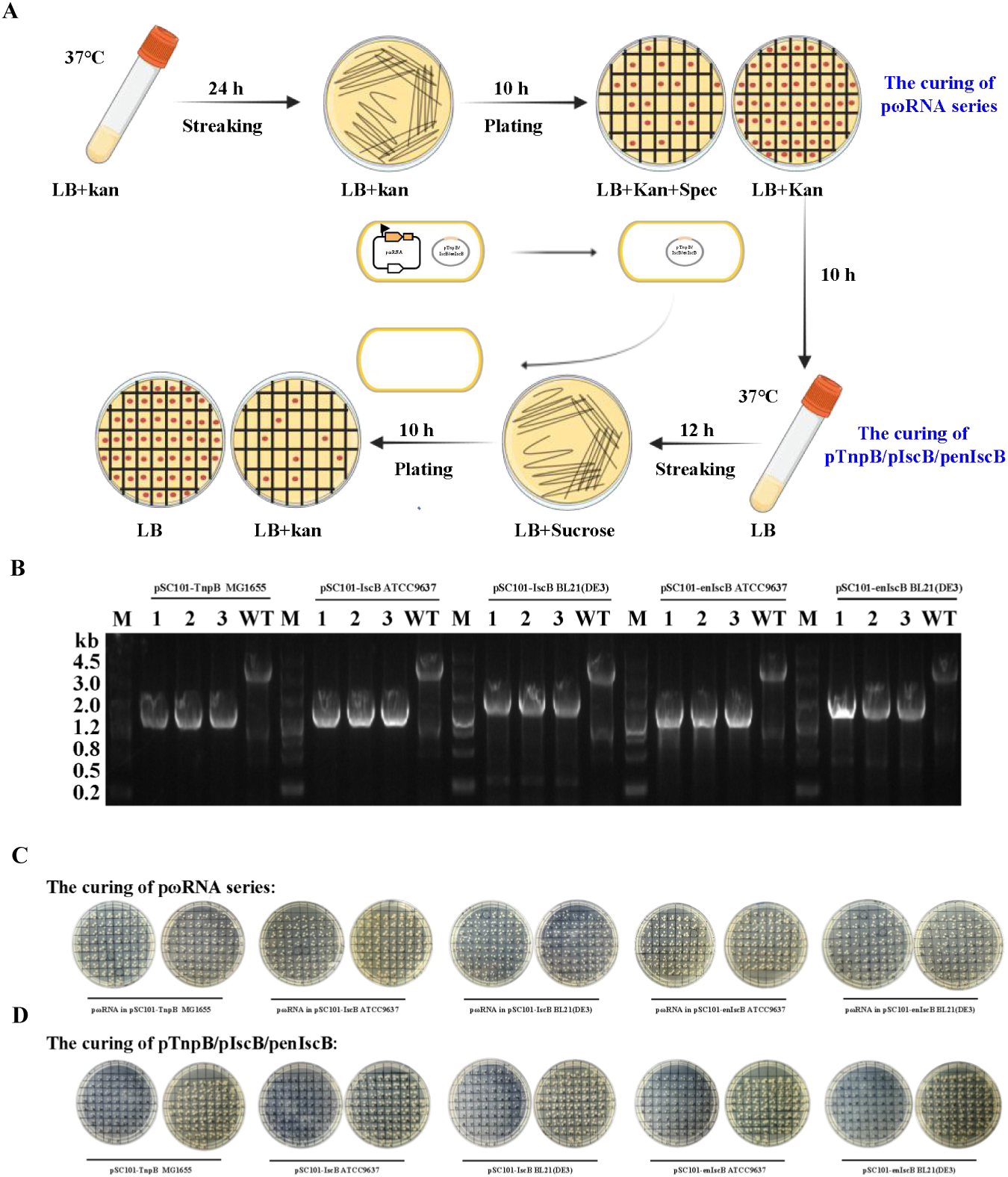
The plasmids curing results of the pTnpB/pωRNA, pIscB/pωRNA and penIscB/pωRNA systems. **A** The two-step plasmid curing process. The pωRNA series was eliminated first, followed by curing the pTnpB/pIscB/penIscB. **B** Reconfirm the genotypes of *E. coli* MG1655Δ*maeB*, *E. coli* ATCC9637Δ*maeB* or *E. coli* BL21(DE3)Δ*maeB.* **C** The results of plasmid curing of the pωRNA series by plating verification. **D** The results of plasmid curing of the pTnpB/pIscB/penIscB by plating verification.

According to the plating results of pωRNA series plasmids curing, the pωRNA-TnpB-*maeB*-THR plasmid curing efficiency was 2/66 in MG1655. While the curing efficiencies of pωRNA-IscB-*maeB* varied from 1/66 to 2/66 in ATCC9637, and the pωRNA-IscB-*maeB* plasmid curing efficiency in BL21(DE3) was similar to the plasmid curing efficiencies in ATCC9637, the efficiency ranging from 1/66 to 3/66 (Fig. 4B, C and Table 1). Similarly, the results of pTnpB/pIscB/penIscB plasmids curing were also analyzed, it was noteworthy that the 66 strains containing the *maeB* mutation were lost pSC101-TnpB in MG1655. Furthermore, the results of pSC101-IscB/pSC101-enIscB plasmids curing elimination in ATCC9637 and BL21(DE3) were similar, the plasmids curing efficiencies all were 66/66 (Fig. 4B, D and Table 1). The above results showed the plasmids elimination of TnpB/IscB/enIscB-based genome editing systems can be accomplished successfully by using two-step plasmid curing strategy.

### Preliminary uncovering the mechanisms of *E. coli* escaping from the editing of TnpB/IscB

Microorganisms may survive under the cleavage of RNA-guided DNA nucleases (called escapers), and the understanding of the causes of the escapers could facilitate the establishment of highly efficient antimicrobial^45^. Therefore, to test the escape rate of *E. coli* under the editing of TnpB/IscB, the *maeB* (non-essential gene) gene were selected as the targeted sites. Testing the escape rate of MG1655 under the cleavage of TnpB was shown as an example, pωRNA-TnpB-*maeB* which targeting *maeB* gene along with the pCK2 plasmid (control group) were respectively transformed into *E. coli* MG1655 containing the pSC101-TnpB expressing TnpB constitutively. Escapers were obtained on LB plates containing kanamycin and spectinomycin, and the escape rate in MG1655 was determined. It was found that the escape rate of *maeB* gene in MG1655 was approximately 10^-4^. And the escape rate in ATCC9637 was also approximately 10^-4^. While in BL21 (DE3), the escape rate was the lowest, 2×10^-5^ (Fig. 5A).

**Fig. 5.**
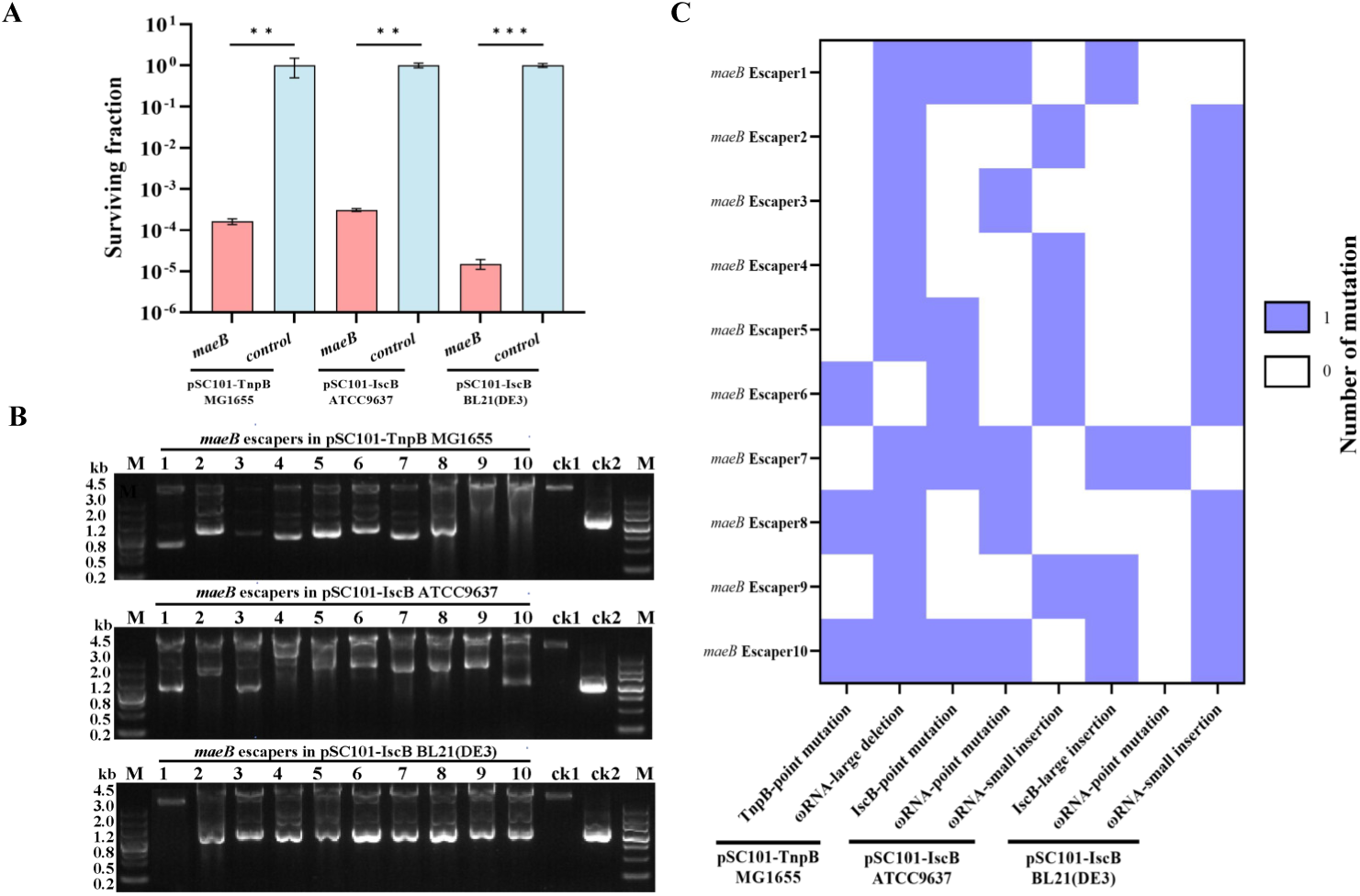
The analysis of escape reason for *E. coli* under the editing of TnpB/IscB. **A** Cell surviving fraction under the targeting of *maeB* site in different *E. coli* strains. **B** The size of pSC101-TnpB/pSC101-IscB and pωRNA series extracted from the escapers obtained at *maeB* site. ck1 were the original pSC101-TnpB/pSC101-IscB, and ck2 was the original pωRNA series. **C** Mutation types and mutation frequency of TnpB, IscB and ωRNA coding sequences from different escapers obtained at *maeB* site. **P < 0.01; ***P < 0.001.

To explore the escaping reason of *E. coli* MG1655, ATCC9637 or BL21 (DE3) under the editing of TnpB/IscB, we subsequently moved to analyze the sizes of plasmids expressing the TnpB/IscB or ωRNA. Ten escapers on the plates were randomly picked, pSC101-TnpB/pSC101-IscB plasmids and pωRNA in those escapers were extracted, and the sizes of those plasmids were checked in an agarose gel along with the controls (ck1 and ck2). Though the change of the size of pSC101-TnpB plasmid was not observed in MG1655, the pωRNA-TnpB-*maeB* plasmid in no. 1, 2, 3, 4, 5, 7, 8, 9, 10 escapers were significantly smaller than the control plasmid (ck2) (Fig. 5B). Meanwhile, the sizes of some pSC101-IscB and pωRNA series plasmids both were changed significantly in ATCC9637 or BL21 (DE3) (Fig. 5B). Next, the DNA coding sequences TnpB, IscB and ωRNA in *E. coli* from 10 escapers obtained at *maeB* site were sequenced respectively. DNA point-mutations were observed in all samples, which including point-mutation in TnpB coding sequence and large-deletion in ωRNA coding sequence in MG1655 (Fig. 5B). While in ATCC9637 and BL21 (DE3), multiple types of mutations were found in all IscB and ωRNA coding sequences, which point-mutation and small-sertion in ωRNA coding sequence dominated the majority (Fig. 5C). The above results showed that point-mutation in ωRNA coding sequence was the main cause of escaper in *E. coli* under the editing of TnpB/IscB. Furthermore, exploring the escaping reasons of *E. coli* escaping from the editing of TnpB/IscB will facilitate the establishment of TnpB/IscB-based genome editing tools and the development of TnpB/IscB-based new antimicrobial drugs.

## Discussion

The growing problem of drug resistance in *E. coli* has been reported in the numerous research, and there is an urgent need to find new strategies to combat drug-resistant bacteria^5–7^. Although the CRISPR-Cas9 has been used for eliminating of specific strains, the excessive size of Cas9 (∼1053-1368 aa) limits its delivery efficiency, which is a bottleneck in the development of the new antimicrobial drugs based on CRISPR-Cas9. Given the recent discoveries of smaller-sized TnpB, IscB and enIscB proteins, and they are considered as the better strategy to combat the drug resistance problem in *E. coli.* In this study, *E. coli* MG1655, ATCC9637 and BL21(DE3) were used as the research objects to test the application of TnpB and IscB in the above strains. First, the toxicity of TnpB, IscB and enIscB proteins in the above three *E. coli* were determined by counting the transformation efficiencies (Supplementary Fig. 1A-D), and it was found that the expression of TnpB protein was toxic to ATCC9637, while the expression of IscB or enIscB proteins were toxic to MG1655. It was speculated that the introduction of TnpB, IscB or enIscB proteins activated the expression of certain genes that were repressed in the host^46^. However, the reason why the expression of TnpB, IscB and enIscB proteins was toxic to those *E. coli* strains has not been well studied.

Subsequently, the cleavage efficiencies of TnpB, IscB and enIscB were further tested in the corresponding nontoxic strains and the transformation efficiencies were counted to determine the cleavage activities in *E. coli* stains (Supplementary Fig. 2, 5). The results showed that TnpB performed the cleavage activity in MG1655, IscB and enIscB could cleave target genes in ATCC9637 and BL21(DE3). Overall, IscB and enIscB proteins exhibited the superior efficiencies for cleaving the *E. coli* genome compared to TnpB protein, and it was speculated that the cleavage activity of TnpB in *E. coli* might be related to the selection of temperature as it was reported that TnpB could performed well with the optimal temperature between 65∼75°C^30^. Furthermore, the testing of gene deletion efficiencies of TnpB, IscB and enIscB in *E. coli* was carried out, and it was found that the TnpB could successfully knock out genes in MG1655, with the editing efficiency of 100%. While in ATCC9637 and BL21(DE3), the deleting of target genes could be achieved by IscB and enIscB, with the efficiency ranging from 13% to 87% (Table 1). These results can fill the gap where TnpB cannot be applied to the gene deletion in ATCC9637 and BL21(DE3).

For the previous research about Cas9, it was found that increasing the copy number of Cas9 could improve the genome editing efficiency in *E. coli*^43^. Therefore, we aimed to optimize these gene editing tools by increasing the copy number of IscB or enIscB. The results showed that increasing the copy number of these protein had little effects on improving of the gene deletion efficiency. Moreover, although the gene deletion in *E. coli* MG1655 can be realized by utilizing TnpB, the homology arm needs to be provided in the form of plasmids. In contrast to IscB and enIscB, the homology arm provided in the form of linear dsDNA greatly makes the editing process more time-efficient and less laborious^47^. More interestingly, it was found that the gene deletion efficiency within *E. coli* could be improved by p15A-enIscB after increasing the content of homology arm from 400 ng to 800 ng, with the deletion efficiency increasing from 0 to 30%. Although the gene deletion efficiencies of the TnpB and IscB systems were similar to the Cas9-based genome editing tools, these smaller-sized DNA nuclease (∼500 aa) confers advantages in terms of the subsequent delivery (Table 1). What’s more, the establishment of TnpB and IscB-based genome editing tools could be optimized by further changing the strength of the promoter, adjusting the length of the homology arm or modifying the guide RNAs^48^. It was found that the pωRNA series plasmids and pTnpB/pIscB/penIscB plasmids can be efficiently eliminated through two-step plasmid curing strategy which taking approximately 4 days for one round curing process (Table 1). It was notewothy that the plasmid curing efficiencies of the TnpB/IscB/enIscB-based genome editing system can accomplish 100%. Finally, we also tested the escaping rate of *E. coli* under the cleavage of TnpB/IscB. It was observed that the escaping reasons of *E. coli* strains escaping from the editing of TnpB/IscB involved point mutation in TnpB, IscB and ωRNA coding sequence, small insertion and large deletion in ωRNA coding sequence. Based on the above preliminary understanding of escaping mechanism could facilitate the applications of TnpB/IscB-based genome editing tools and the development of new antimicrobial drugs.

In summary, we took the lead to systematically characterize the application of TnpB, IscB and enIscB in *E. coli* strains. On the one hand, establishing TnpB-based genome editing tool in *E. coli* MG1655 successfully. On the other hand, it was found that the genome editing in *E. coli* ATCC9637 and BL21(DE3) can be achieved by using IscB and enIscB. Meanwhile, the TnpB, IscB and enIscB editing plasmids were successfully cured. Furthermore, we also analyzed how *E. coli* escapes the editing of TnpB/IscB. Totally, due to the smaller size of TnpB, IscB and enIscB compared to Cas9, they have more advantages and will provide new research ideas and methods for the eliminating of drug-resistant microorganisms.

## Materials and methods

### Bacterial strains and culture methods

*E. coli* DH5α was used as the cloning host to construct plasmids. *E. coli* MG1655, BL21(DE3) and ATCC9637 strains were used in this study. These strains were cultured in LB medium (0.5% yeast extract, 1% [w/v] tryptone, 1% [w/v] NaCl) at 37°C. Antibiotic spectinomycin (100 μg/mL), kanamycin (50 μg/mL) and chloramphenicol (25 μg/mL) were supplemented into LB medium when required. Strains were stored at -80°C at a final concentration of 20% glycerol, and the strain recovery was carried out by streaking the culture on LB plate and inoculating the isolated colony in LB broth. All strains are listed in Supplementary Table 1.

### Reagents and enzymes

Restriction endonucleases used in this study were purchased from Thermo Fisher Scientific (Waltham, MA, USA). DNA polymerases, including 2×Phanta Flash Master Mix (Vazyme Biotechnology Co., Ltd., Nanjing, China) for high-fidelity DNA amplification and 2×Es Taq MasterMix (Dye) (Jiangsu Cowin Biotech Co., Ltd., Taizhou, China) for colony PCR. Reagent kit for plasmid extraction was obtained from Tiangen (Tiangen Biotech Co., Ltd., Beijing, China), and those for DNA purification was sourced from Transgen (Beijing, China), following the manufacturer’s instructions. The ClonExpress One Step Cloning Kit (Vazyme Biotechnology Co., Ltd., Nanjing, China) was used for the assembly of plasmids.

### Plasmids construction

All plasmids and primers used in this study are listed in Supplementary Table 1, 2. *E. coli* DH5α was used for the maintenance and construction of plasmids. The TnpB gene sequence was derived from the plasmid pSisTnpB-ωRNA^30^.The IscB and enIscB gene sequences were obtained from the pIscB-ωRNA and penIscB-ωRNA plasmids respectively^36^. Take the pSC101-TnpB plasmid as an example, the TnpB gene sequence was assembled into linearized pSC101 (∼5 copies) plasmid vector to generate pSC101-TnpB plasmid. The constructions of pSC101-IscB/p15A-IscB or pSC101-enIscB/p15A-enIscB plasmids were similar to the construction of pSC101-TnpB (The TnpB, IscB and enIscB proteins were expressed constitutively in these above plasmids).

For the construction of pωRNA-TnpB-targetX (targetX was represented as the target gene) series plasmids, the gRNA sequence on the original pEcgRNA plasmid was replaced with the ωRNA sequence derived from pSisTnpB-ωRNA plasmid^30^. Similarly, for the construction of pωRNA-IscB-targetX series plasmids, the gRNA on the pEcgRNA was replaced with the ωRNA sequence derived from pIscB-ωRNA plasmid^36^. While for pωRNA-TnpB-targetX-THR plasmids, the homology arms (∼500 bp) amplified from the genomic DNA of *E. coli* were inserted into pωRNA-TnpB-targetX to generate the plasmids pωRNA-TnpB-targetX-THR.

### Plasmid transformation and testing the toxicity of TnpB, IscB and enIscB in *E. coli*

The competent cells for *E. coli* (including MG1655, ATCC9637, and BL21(DE3)) electroporation were prepared as the method mentioned previously^40,49^. Taking the toxicity testing of TnpB for example, 100 μL of competent cells of MG1655, ATCC9637, and BL21(DE3) was mixed with 100 ng TnpB (including pSC101-TnpB and p15A-TnpB) respectively. The pCK1 plasmid (∼10 kb) which was nontoxic to *E. coli* MG1655, ATCC9637, and BL21(DE3) was used as the control. The mixture was electroporated in a cooled 1-mm Gene Pulser cuvette (Bio-Rad, Hurcules, USA) at 1.8 kV. The electroporated mixture was immediately suspended in 1 mL of fresh LB broth. Incubating the cells at 37°C for 1 h and plating the recovered cells on LB solid medium containing kanamycin (for pSC101-TnpB) or chloramphenicol (for p15A-TnpB). Then incubating the plates overnight at 37°C. The colonies grown on plates were counted as colony-forming units (CFU), and the plasmid transformation efficiency (CFU/μg plasmid DNA) was calculated. If the transformation efficiency of the TnpB was significantly lower than the control plasmid, the protein was recognized as toxic to the host. While, if the transformation efficiency of TnpB was equivalent to the control plasmid, the protein was recognized as nontoxic to the host.

For testing the toxicity of IscB and enIscB in *E. coli* strains, the steps were similar to the above method. Plasmids that carrying IscB or enIscB proteins (including pSC101-IscB/p15A-IscB, pSC101-enIscB/p15A-enIscB) were transformed into *E. coli* respectively, and the transformation efficiencies were counted to determine the toxicity in *E. coli* stains.

### Testing the cleavage activities of TnpB, IscB and enIscB in *E. coli*

The testing of cleavage activities was performed by using the proteins that were nontoxic to the corresponding hosts. The cleavage testing of TnpB was shown as an example, 100 µL of competent cells of *E. coli* carrying plasmids that expressing TnpB (including pSC101-TnpB/p15A-TnpB) was mixed with 100 ng of pωRNA-TnpB-targetX series plasmid and the pCK2 control plasmid (∼2.5 kb) that did not target any sites in the genome respectively. The mixture was suspended in 1 mL of fresh LB broth after electroporation. Incubating the cells at 37°C for 1 h and plating the recovered cells on LB solid medium containing spectinomycin and kanamycin (for pSC101-TnpB) or spectinomycin and chloramphenicol (for p15A-TnpB). The plates were then incubated overnight at 37°C. The CFU were counted to calculate the plasmid transformation efficiency. If the transformation efficiency of TnpB was similar with the control plasmid, it was considered that the protein could not cleave the host genomic DNA. While, if the transformation efficiency of the TnpB was significantly lower (at least two orders of magnitude) than the control plasmid, it was considered that the protein could cleave the host genomic DNA.

The cleavage testings of IscB and enIscB were similar with the above process. Plasmids that carrying IscB or enIscB proteins (including pSC101-IscB/p15A-IscB, pSC101-enIscB/p15A-enIscB) were transformed into *E. coli* respectively, and the transformation efficiencies were counted to determine the cleavage activities in *E. coli* stains.

### Verifying the gene deletion efficiencies of TnpB, IscB and enIscB in *E. coli*

The testing of gene deletion efficiency of TnpB was shown as an example, 10 mM of arabinose was added to the culture of *E. coli* carrying the pSC101-TnpB or p15A-TnpB to induce the expression of the λ-Red system respectively. Competent cells for electroporation were prepared subsequently. 100 µL of competent cells was mixed with 100 ng of pωRNA-TnpB-targetX-THR plasmid. The mixture was electroporated and immediately suspended in 1 mL of fresh LB broth. Next, the cells were incubated for 1 h at 37℃, and plating the recovered cells on the LB solid medium containing spectinomycin and kanamycin (for pSC101-TnpB) or spectinomycin and chloramphenicol (for p15A-TnpB), then incubating the plates overnight at 37℃. Then, about 15 colonies were randomly selected and verified by colony PCR, the primers complementary to the sequences ∼50 bp upstream and downstream of the homologous arms on the genome were used. The PCR products were subsequently sequenced to verify the positive colonies, and the corresponding wild-type strains were used as controls.

The verification of gene deletion efficiency of IscB or EnIscB (including pSC101-IscB/p15A-IscB, pSC101-enIscB/p15A-enIscB) was similar to TnpB, the only difference was that homology arms (∼400 ng) were provided in the form of linear dsDNA, rather than being placed on plasmids.

### Plasmid curing

For the curing of pωRNA series plasmid, the pωRNA-TnpB-targetX-THR was taken as an example. The edited colony harboring both pSC101-TnpB and pωRNA-TnpB-targetX-THR plasmid was inoculated into 5 mL of LB medium containing kanamycin (50 µg/mL), the culture was incubated overnight with shaking at 220 rpm, and spread onto LB medium containing kanamycin (50 µg/mL). The colonies that grew on kanamycin LB plates after incubation overnight at 37℃ were randomly picked and screened on LB plates carrying kanamycin (50 µg/mL) and spectinomycin (50 µg/mL). The colonies were confirmed as cured by determining their sensitivity to spectinomycin. The curing processes of pωRNA-IscB-targetX plasmids were similar to the plasmid curing of pωRNA-TnpB-targetX-THR.

For curing the pTnpB/pIscB/penIscB series plasmids, the pSC101-TnpB was taken as an example. The colonies cured of plasmid pωRNA-TnpB-targetX-THR were further inoculated in liquid LB medium at 37℃ and incubated overnight with shaking at 220 rpm. Approximately 10 μL of the cells was then spread on LB plates containing sucrose (10 g/L) and incubated overnight at 37℃. Individual colonies were then randomly picked and screened on LB plates with and without kanamycin (50 µg/mL). The colonies sensitive to kanamycin were cured of pSC101-TnpB. Similarly, the curing processes of pSC101-IscB/pSC101-enIscB plasmids were similar to the plasmid curing of pSC101-TnpB.

### Workflow of verifying the reason of the escape

After obtaining the escapers on plates, about 10 escapers were randomly selected for *maeB* site, the pSC101-TnpB/pSC101-IscB and pωRNA series plasmids of the escapers were extracted, and their sizes were checked by agarose gel electrophoresis. In the case of the size of the plasmids showing obvious abnormality, DNA coding sequencing of the key elements (including the TnpB, IscB and ωRNA coding sequencing) of the plasmid was performed to reveal the specific mutation.

## Acknowledgements

This study was supported by Sichuan Science and Technology Program (Number: 2024NSFSC0373), Luzhou Laojiao Co., Ltd (Number: 2022HX07). We would like to thank Prof. Peng Nan for providing the pSisTnpB-ωRNA related plasmids; Prof. Hui Yang and Yingsi Zhou for providing the pIscB/penIscB-ωRNA related plasmids.

## Author contributions

Q.L., H.J.T., and J.G., conceived the project and designed the experiments. H.J.T., and J.G. and M.J.S., carried out the experiments. H.J.T., and J.G., analyzed the data. Q.L., H.J.T., and J.G., wrote the manuscript. All authors discussed the results and contributed to the final manuscript.

