## Supplementary Table 1-2. Supplementary Fig. 1-6 for "Characterization of the genome editing with miniature nucleases TnpB, IscB and enIscB in *Escherichia coli* strains": supplementary.pdf

**Supplementary Figure 1-6**

**Supplementary Table 1. Plasmids and strains used in this study.**

| Strains or plasmids | Description | Source |
| --- | --- | --- |
| Strains | | |
| *E. coli* DH5α | Commercial transformation host | GIBCO BRL, life Technologies |
| *E. coli* MG1655 | Host for testing the genome editing efficiency | Lab storage |
| *E. coli* ATCC9637 | Host for testing the genome editing efficiency | Lab storage |
| *E. coli* BL21(DE3) | Host for testing the genome editing efficiency | Lab storage |
| Plasmids | | |
| pSC101-TnpB | Derived from pSC101, constitutively expressing the TnpB | This study |
| p15A-TnpB | Derived from p15A, constitutively expressing the TnpB | This study |
| pSC101-IscB | Derived from pSC101, constitutively expressing the IscB | This study |
| p15A-IscB | Derived from p15A, constitutively expressing the IscB | This study |
| pSC101-enIscB | Derived from pSC101, constitutively expressing the enIscB | This study |
| p15A-enIscB | Derived from p15A, constitutively expressing the enIscB | This study |
| pωRNA-TnpB-*maeB* | Derived from pSisTnpB-ωRNA, target *maeB* gene in MG1655 | This study |
| pωRNA-TnpB-*umuDC* | Derived from pSisTnpB-ωRNA, target *umuDC* gene in MG1655 | This study |
| pωRNA-TnpB-*maeB*-THR | Derived from pSisTnpB-ωRNA, carrying the homology arms of *maeB* gene for repairing the DSB in MG1655 | This study |
| pωRNA-TnpB-*umuDC*-THR | Derived from pSisTnpB-ωRNA, carrying the homology arms of *umuDC* gene for repairing the DSB in MG1655 | This study |
| pωRNA-IscB-*rpoS* | Derived from pIscB-ωRNA, targeting *rpoS* gene in ATCC9637 and BL21(DE3) | This study |
| pωRNA-IscB -*maeB* | Derived from pIscB-ωRNA, targeting *maeB* gene in ATCC9637 and BL21(DE3) | This study |
| pωRNA-IscB -*umuDC* | Derived from pIscB-ωRNA, targeting *umuDC* gene in ATCC9637 and BL21(DE3) | This study |

**Supplementary Table 2. Oligonucleotides used in this study.**

| **Oligos** | **Sequence (5'→3')** |
|---|---|
| pSC101-TnpB-up | ttttattttaggaggcaaaaatgcatcaccatcaccacca |
| pSC101-TnpB-dn | atcttcatctaaaatatacttcacgatggagatttcctgc |
| pSC101-arac-up | gcaggaaatctccatcgtgaagtatattttagatgaagat |
| pSC101-arac-dn | actggtattggcacaaacctgattcc |
| pSC101-rha-up | ggaatcaggtttgtgccaataccagt |
| pSC101-rha-dn | tggtggtgatggtgatgcatttttgcctcctaaaataaaa |
| TnpB-verf-up | ctttccctgcagctgaac |
| TnpB-verf-dn | tatgtatccgctcatgtc |
| TnpB-ce-1 | ctttccctgcagctgaac |
| TnpB-ce-2 | gtagaacaactgttcaccg |
| red-ce-1 | gtgtgtttgtatgccctga |
| red-ce-2 | atgatttgcccaaacaggtc |
| red-ce-3 | attcactaacccccctttcct |
| red-ce-4 | cgaaggcagagaaatcacgg |
| red-ce-5 | gtggtttcgaggccataaag |
| pωRNA-TnpB-F | gtcctaggtataatactagtttaagaaggacttgactttg |
| pωRNA-TnpB-R | ctctagagaattcaaaaaaacaaaaaacccctcaagaccc |
| pωRNA-TnpB-ptF | gggtcttgaggggttttttgtttttttgaattctctagag |
| pωRNA-TnpB-ptR | caaagtcaagtccttcttaaactagtattatacctaggac |
| pωRNA-Tverf-up | ggatccttgacagctagctc |
| pωRNA-Tverf-dn | cttatcatccccttttgctt |
| pωRNA-ce-up | catgttctttcctgcgttat |
| pωRNA-TnpB-*maeB* | gtaagggcggggttgttcactgtcgtgctcgaaagaacgggtcgacggccggcatggtc |
| pωRNA-TnpB-*umuDC* | gtaagggcggggttgttcacagcgcggatcggattcgttagtcgacggccggcatggtc |
| pωRNA-dn-univers | gtgaacaaccccgcccttac |
| *umuDC*-pωRNA-up | ttccagtgccagagcagagagtcgacctgcagaag |
| *umuDC*-pωRNA-dn | tcgctttaaacaggagtattcacagagaattcaaaaaaacaa |
| *umuDC*-ωHR-upup | gtgaatactcctgtttaaagcga |

| | |
|---|---|
| *umuDC*-ωHR-updn | ctttttcctgccgctatataataatctgcctgaagttat |
| *umuDC*-ωHR-dnup | ataacttcaggcagattattatatagcggcaggaaaaag |
| *umuDC*-ωHR-dndn | ctgctctggcactggaa |
| *maeB*-pωRNA-up | gacccaggaacaacaggtggtcagagtcgacctgcagaag |
| *maeB*-pωRNA-dn | cgctgatgtgcatatcggttagagaattcaaaaaaacaaaaaacc |
| *maeB*-ωHR-upup | aaccgatatgcacatcag |
| *maeB*-ωHR-updn | tacgtgaaaggaacaaccaatttttttaactctcacgct |
| *maeB*-ωHR-dnup | agcgtgagagttaaaaaaaattggttgttcctttcacgta |
| *maeB*-ωHR-dndn | gaccacctgttgttcctg |
| pωHR-verf-up | aggaaatctccatcgtgagg |
| pωHR-verf-dn | cggagccgtacaaatgtacg |
| pSC101/p15A-IscB-up | ctttttattttaggaggcaaaaatgatggccgtggtgtacgtga |
| pSC101/p15A-IscB-dn | gaaataatcttcatctaaaatatactttacacgtagatctgcaggccgccattgtttct |
| pSC101/p15A-pt-up | agaaacaatggcggcctgcagatctacgtgtaaagtatattttagatgaagattatttc |
| pSC101/p15A-pt-dn | tcacgtacaccacggccatcatttttgcctcctaaaataaaaag |
| IscB-verf-up | ctttccctgcagctgaac |
| IscB-verf-dn | tatgtatccgctcatgtc |
| IscB-ce-1 | atcgcgagcccatttatacc |
| IscB-ce-2 | ctaagctgatgaaagacaga |
| IscB-ce-3 | ccgtcaagttgtcataataa |
| pωRNA-IscB-F | gtcctaggtataatactagtagtcttccgaagacttggc |
| pωRNA-IscB-R | ctctagagaattcaaaaaaaccgttcctccttttgtaaaa |
| pωRNA-IscB-ptF | ttttacaaaaggaggaacggtttttttgaattctctagag |
| pωRNA-IscB-ptR | actagtattatacctaggactgagctagctgtcaa |
| pωRNA-Iverf-up | ggatccttgacagctagctc |
| pωRNA-Iverf-dn | cttatcatccccttttgctt |
| pωRNA-ce-dn | cttatggagctgcacatga |
| pωRNA-IscB-*rpoS* | aggtataatactagtaaaggccttagtagaaggctcttccaactttatggttgcgaccg |
| pωRNA-IscB-*maeB* | aggtataatactagtcccgaaagtgctgaccggctcgtccaactgcggttgaacgagca |
| pωRNA-IscB-*umuDC* | aggtataatactagtgtttgcaccgacgaagggctcgtccaactgcggttgaacgagca |
| pωRNA-dn-univers | actagtattatacctaggactgagctagctgtcaa |
| *rpoS*-F-MG | gaacagagtgctaacaaaat |
| *rpoS*-R-MG | agttacgacagctttttcag |

| | |
|---|---|
| *rpoS*-R-AT | agttacgacagctttttcag |
| *maeB*-F | aaccgatatgcacatcagc |
| *maeB*-R | gaccacctgttgttcctg |
| *umuDC*-F | tatgttgtttatcaagcctg |
| *umuDC*-R | atttactgagggtcaaataa |
| *rpoS*-ko-up | aaaattccaccgttgctgtt |
| *rpoS*-ko-dn | aaccgctatgatagcgac |
| *maeB*-ko-up | tgataacgcttcttctactg |
| *maeB*-ko-dn | aacttctgttgacacgcga |
| *umuDC*-ko-up | cataagtaaggttttaatat |
| *umuDC*-ko-dn | ttgaaccaatgttcacaagg |

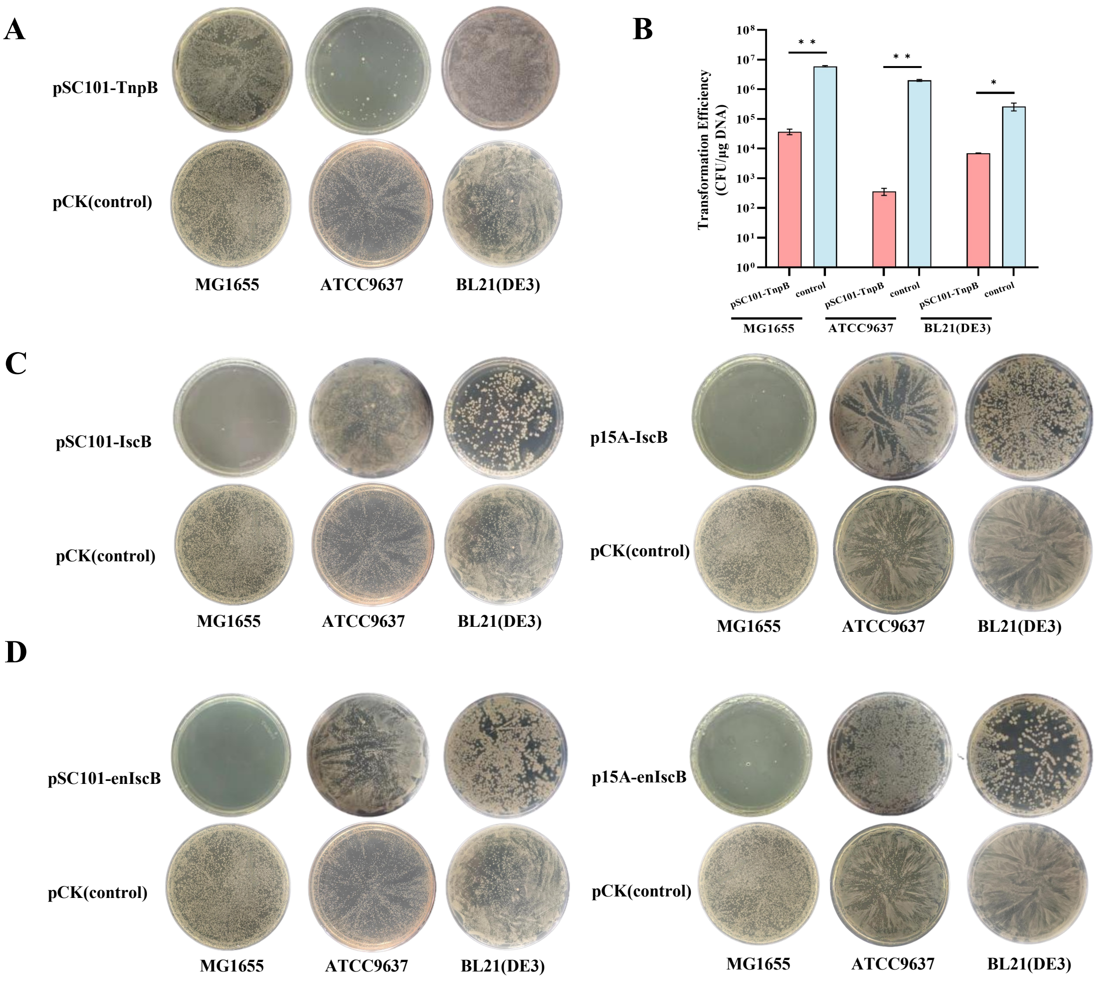


**Supplementary Fig. 1 The toxicity testings of TnpB, IscB and enIscB in *E. coli* MG1655, ATCC9637 and BL21(DE3).**

(A) The verification results of toxicity of pSC101-TnpB in *E. coli* MG1655, ATCC9637 and BL21(DE3). (B) The plating results for the verification of toxicity of p15A-TnpB in *E. coli* MG1655, ATCC9637 and BL21(DE3) (C) The verification results of toxicity of pSC101-IscB and p15A-IscB in *E. coli* MG1655, ATCC9637 and BL21(DE3). (D) The verification results of toxicity of pSC101-enIscB and p15A-enIscB in *E. coli* MG1655, ATCC9637 and BL21(DE3).

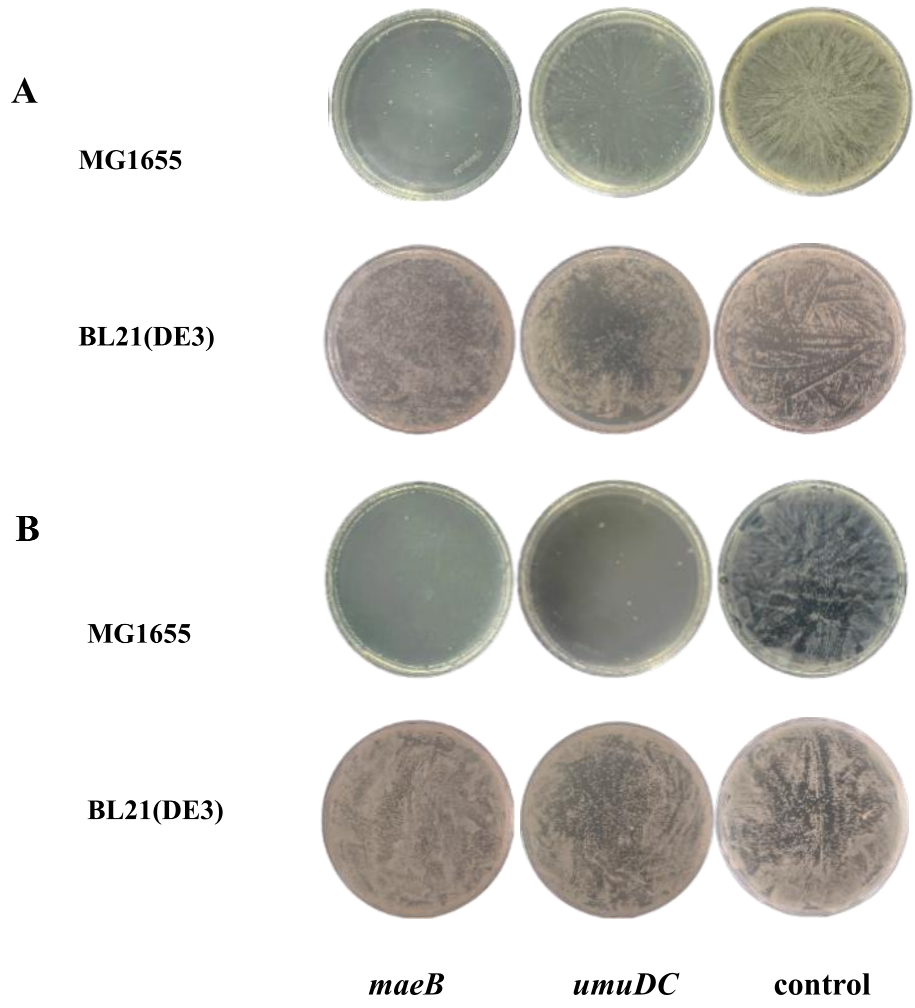


**Supplementary Fig. 2 The testings of cleavage activity of TnpB in *E. coli* MG1655 and BL21(DE3).**
(A) The genomic cleavage activity in MG1655 and BL21(DE3) by using pSC101-TnpB. (B) The genomic cleavage activity in MG1655 and BL21(DE3) by using p15A-TnpB.

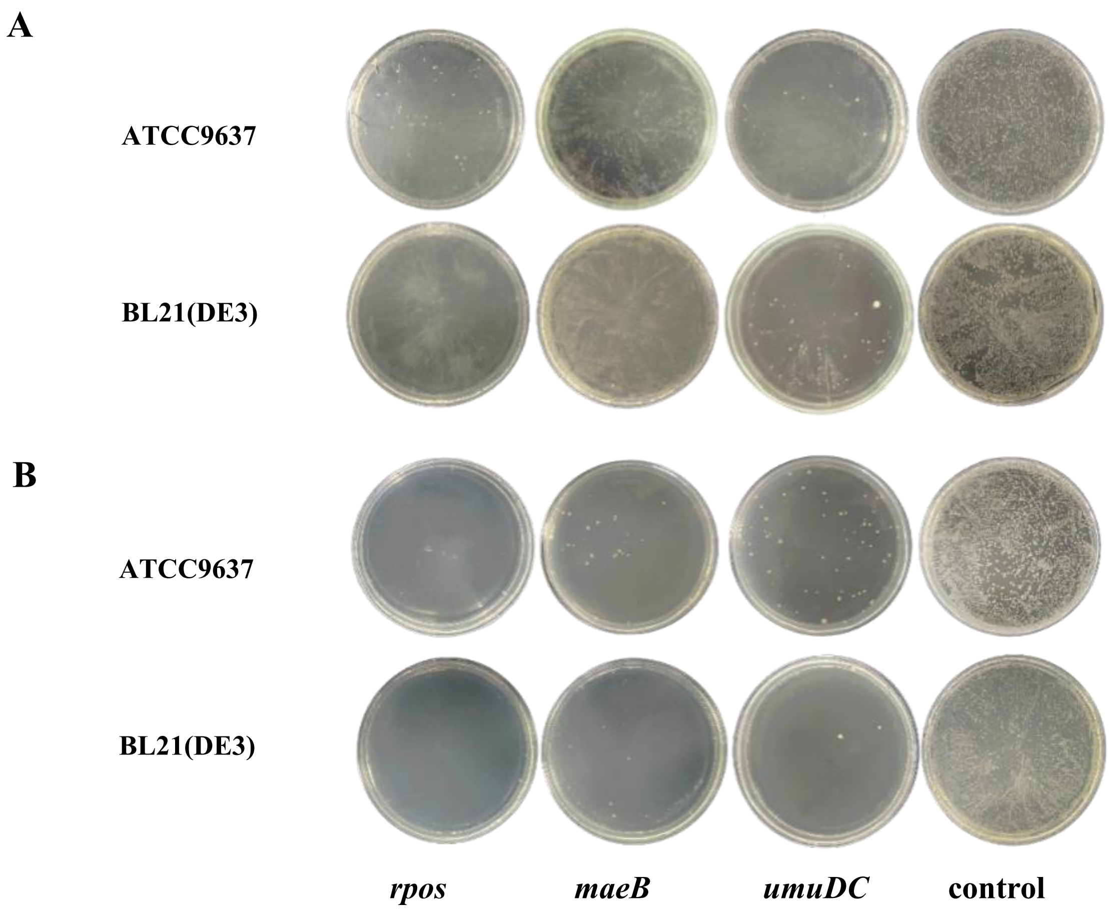


**Supplementary Fig. 3 The testings of cleavage activity of IscB in *E. coli* ATCC9637 and BL21(DE3).** (A) The genomic cleavage activity in ATCC9637 and BL21(DE3) through using pSC101-IscB. (B) The genomic cleavage activity in ATCC9637 and BL21(DE3) through using p15A-IscB.

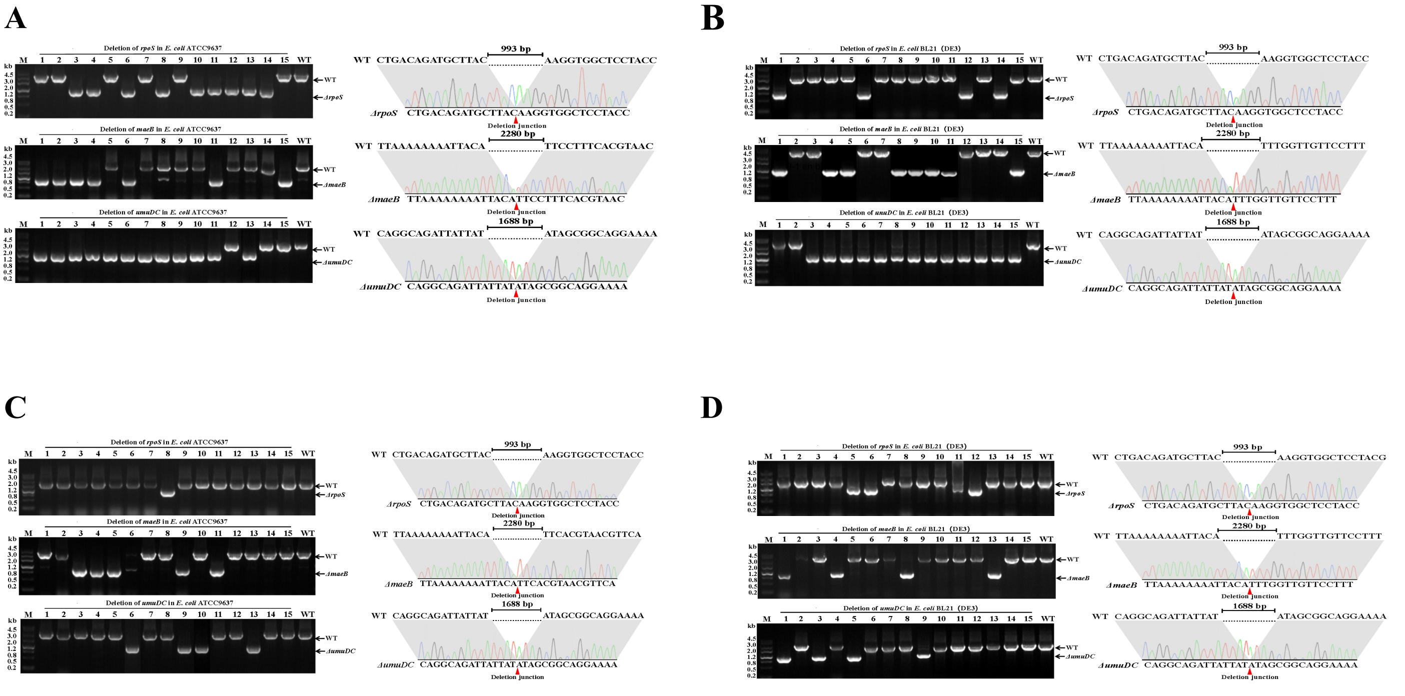


**Supplementary Fig. 4 The genome editing could be achieved in *E. coli* ATCC9637 and BL21(DE3) by using IscB.**

(A) The colony PCR and DNA sequencing results of the deletion of *rpoS, maeB* and *umuDC* genes in ATCC9637 by using pSC101-IscB. (B) The colony PCR and DNA sequencing results of the deletion of *rpoS, maeB* and *umuDC* genes in BL21(DE3) by utilizing pSC101-IscB. (C)The colony PCR and DNA sequencing results of the deletion of *rpoS, maeB* and *umuDC* genes in ATCC9637 by using p15A-IscB. (D) The colony PCR and DNA sequencing results of the deletion of *rpoS, maeB* and *umuDC* genes in BL21(DE3) by utilizing p15A-IscB.

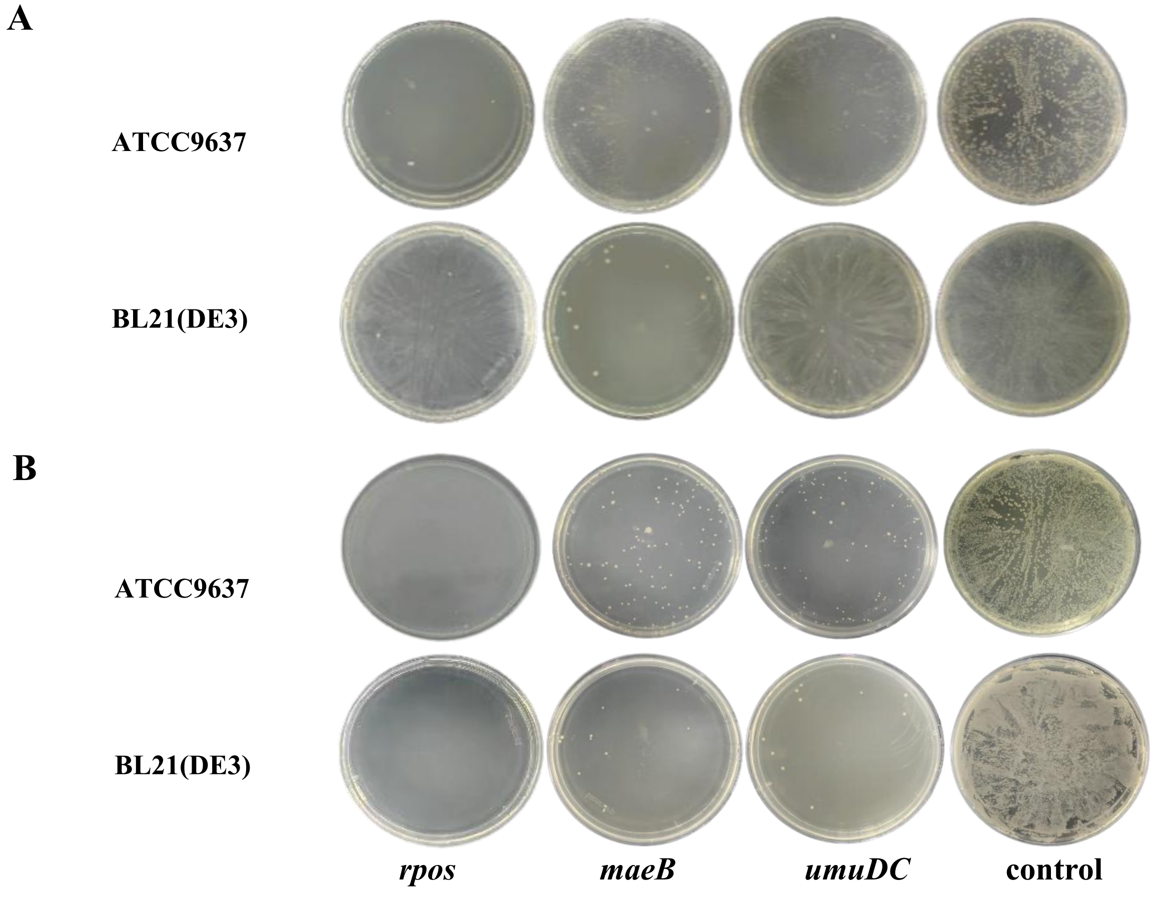


**Supplementary Fig. 5 The testings of cleavage activity of enIscB in *E. coli* ATCC9637 and BL21(DE3).** (A) The genomic cleavage activity in MG1655 and BL21(DE3) by using pSC101-enIscB. (B) The genomic cleavage activity in MG1655 and BL21(DE3) by using p15A-enIscB.

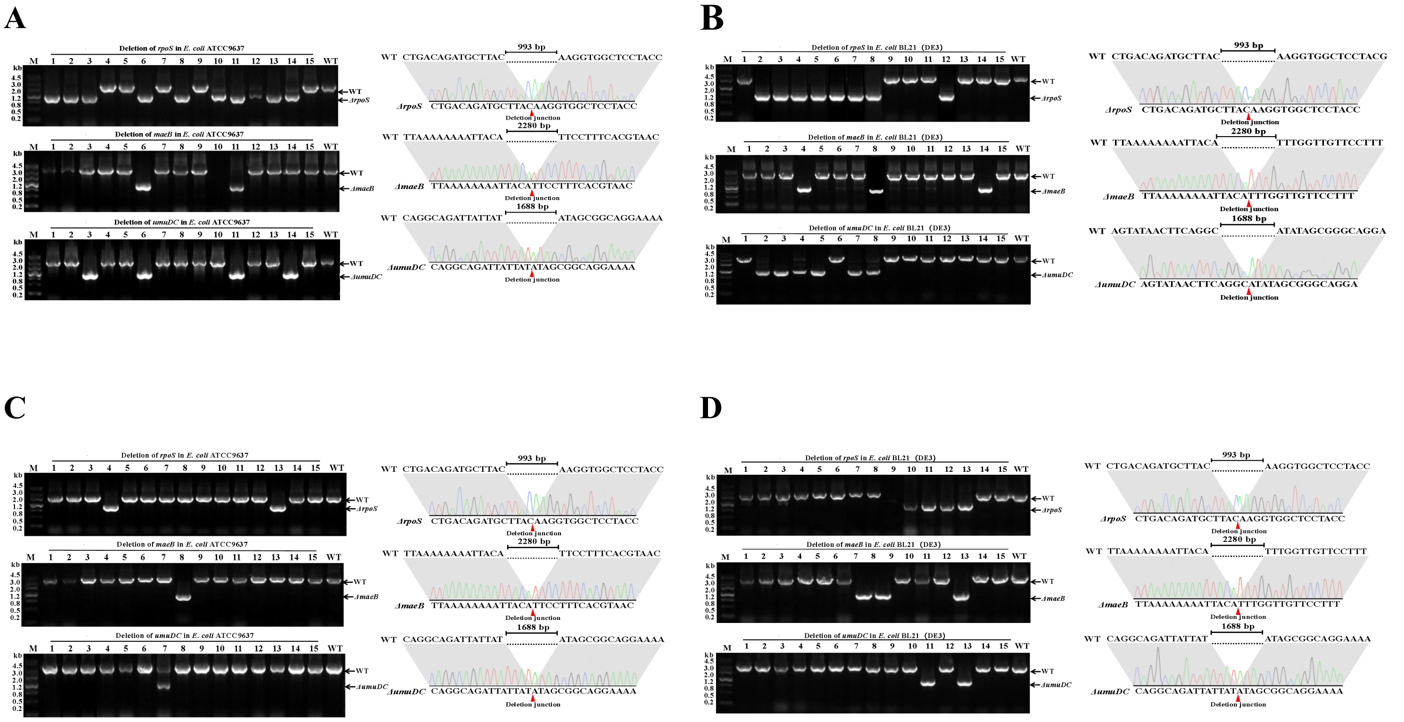

**Supplementary Fig. 6 The genome editing could be achieved in *E. coli* ATCC9637 and BL21(DE3) by using enIscB.**

(A) The colony PCR and DNA sequencing results of the deletion of *rpoS, maeB* and *umuDC* genes in ATCC9637 by using pSC101-enIscB. (B) The colony PCR and DNA sequencing results of the deletion of *rpoS, maeB* and *umuDC* genes in BL21(DE3) by utilizing pSC101-enIscB. (C)The colony PCR and DNA sequencing results of the deletion of *rpoS, maeB* and *umuDC* genes in ATCC9637 by using p15A-enIscB. (D) The colony PCR and DNA sequencing results of the deletion of *rpoS, maeB* and *umuDC* genes in BL21(DE3) by utilizing p15A-enIscB.
